# SUMO E2/E3 Enzyme Co-Recognize Substrates and Confers High Substrates Specificity

**DOI:** 10.1101/2021.08.16.456439

**Authors:** Zhehao Xiong, Ling Jiang, Yan Liu, Jun Li, Jiayu Liao

## Abstract

Although various technologies can determine protein-protein interaction affinity, *K_d_*, current approaches require at least one interacting protein to be purified. Here, we present a development of high-throughput approach to determine protein interaction affinity for un-purified interacting proteins using quantitative FRET assay. We developed this approach first using SUMO E2 conjugating enzyme, Ubc9, and SUMO substrate, RanGap1c. The interaction affinities from two purified proteins and two unpurified proteins in the presence of BSA, bacterial extracts or two mixtures are all in excellent agreement with that obtained from the SPR measurement. We then applied this approach to determine the interaction affinities of SUMO E3 PIAS1 and Ubc9 with a SUMO substrate, influenza virus protein, NS1, and, for the first time, the SUMO E3 ligase-substrate interaction affinity is determined, which enables us to provide a kinetics explanation for the two-enzyme substrate recognition mode. In general, our studies allow high-throughput determination of protein interacting affinity without any purification, and both the approach and scientific discoveries can be applied to proteins that are difficult to be expressed in general.

- Protein interaction dissociation constant, K_d_, determination without any protein purification;
- Co-recognition of substrates by two enzymes in one reaction;
- Potential affinity improvement mechanism of substrate recognition revealed with kinetics parameters *in vivo*.

## INTRODUCTION

Protein-protein interaction plays central roles in all the physiological and its alternation many pathological processes(Braun & Gingras, 2012; Nibbe *et al*, 2011). Various approaches have been developed to determine protein interaction affinity, such as Surface plasmon resonance (SPR) and Calorimetric methods (for example, ITC-isothermal titration calorimetry and DSC-differential scanning calorimetry, radioactive labeling binding assay, ultracentrifugation and fluorescence polarization (FP)(Zhou *et al*, 2016). These methods have greatly expanded the understanding of protein functions and dynamics ranging from ligand-receptor binding to signal transductions. However, these methods usually require tedious procedures and/or expensive instruments. Most importantly, current approaches for protein interaction affinity determinations require at least one protein partner to be purified, and a relatively large amount of protein is needed in most approaches. Although several approaches, including tandem affinity purification with mass spectrometry (TAP-MS) and yeast two-hybrid (Y2H), have been developed to determine protein complex in the cells, due to those technique challenges, affinities for the difficult-to-be expressed proteins and kinetics in large-scale proteomics network are still largely unknown, and the consequent functional capabilities of those proteins in physiology and pathology remain poorly understood(Huttlin *et al*, 2017; Krogan *et al*, 2006; Rigaut *et al*, 1999; Syafrizayanti *et al*, 2014; Uetz *et al*, 2000).

Quantitative FRET approaches, such as quantitative FRET imaging and biochemical quantitative FRET assay, use quantitative three-cube or FRET analysis or titration ratio-metric FRET assay to obtain FRET signals corresponding to the protein interaction, which, in theory, could be used to determine protein interaction affinity *K_d_* in a mixture(Gordon *et al*, 1998). However, the quantitative three-cube approach requires measurements of molar extinction coefficients of fluorophores and FRET efficiency. It also needs estimates of FRET efficiency and many instrument-dependent parameters during measurements, which makes it difficult to turn the approach into a general methodology(Erickson *et al*, 2001; Erickson *et al*, 2003). The FRET analysis approach uses a point-to-point subtraction method to measure FRET signal and fluorescent ratio of emission of acceptor and donor wavelength, which does not exclude signals of direct emissions of donor and acceptor(Martin *et al*, 2008; Mehta *et al*, 2009). The application of titration ratio-metric FRET assay to determine protein interaction affinity usually yields *K_d_* values larger than those determined by SPR or ITC. The lack of accuracy and robustness of these approaches for *K_d_* determinations in a mixture is due to multiple fluorescence parameter estimations and difficulty of absolute FRET signal determination.

A recent quantitative FRET (qFRET) approach determines the absolute FRET signal from the subtraction of the total fluorescence signal with the fluorescent signals of free donor and free acceptor obtained from the donor or acceptor emissions multiplied by their cross-wavelength correlation constants(Liao *et al*, 2015; Song *et al*, 2011). The *K_d_* value determined with this approach, which we refer to as qFRET, is in an excellent agreement with the values determined using SPR and ITC, demonstrating its accuracy. In addition, this approach was carried out in the multiwell plate and therefore it can be expanded to the high-throughput mode. However, its potential for determination of protein interaction affinity without purification has not been explored and this represents an important application direction.

Here, we apply quantitative FRET analysis method, for the first time, to determine protein interaction affinities of interactive partners from cell extract mixtures without any purification in a high-throughput setting. We also determine the protein interaction affinities of SUMOylation E3 ligase, PIAS1, to E2 conjugating enzyme, UBC9 and SUMOylation substrate, influenza virus protein NS1, to provide kinetics basis for roles that SUMO E3 ligase in the SUMO conjugation process.

## MATERIALS AND METHODS

### DNA constructs

The open reading frames of CyPet and YPet were amplified by PCR with primers containing NheI-SalI sites. The size of PCR products were 729 and 729 bp, respectively. RanGAP1c and Ubc9 were amplified by PCR with primers containing SalI-NotI sites. All these four genes were cloned into pCRII-TOPO vector (Invitrogen). Then the fragments encoding RanGAP1c and Ubc9 were extracted after a digestion by SalI-NotI and inserted into pCRII-CyPet or pCRII-YPet linearized by SalI and NotI. After the sequences were confirmed by sequencing, the cDNAs encoding CyPetRanGAP1c and YPetUbc9 were cloned into the NheI-NotI sites of pET28(b) vector (Novagen).

### Protein expression, purification and concentration determination

BL21(DE3) *Escherichia coli* cells were transformed with pET28(b) vectors encoding CyPetRanGAP1c or YPetUbc9. The transformed bacteria were plated on LB agar plates containing 50 μg/mL kanamycin, and single colonies were picked up and inoculated in 2xYT medium to an optical density at 600 nm of 0.5–0.8. The expression of polyhistidine-tagged recombinant proteins was induced with 0.2 mM isopropyl β-D-thiogalactoside. Bacterial cells were collected by centrifugation at 6,000 rpm 10 min, resuspended in binding buffer (20 mM Tris-HCl, pH 7.4, 500 mM NaCl and 5 mM imidazole), and sonicated with an ultrasonic liquid processor (Misonix). Cell lysates containing recombinant proteins were cleared by centrifugation at 35,000 g for 30 min. The polyhistidine-tagged recombinant proteins were then purified from bacterial lysates with Ni^2+^-NTA agarose beads (QIAGEN) and washed by three different washing buffers (Washing buffer 1 contained 20 mM Tris-HCl, pH 7.4, 300 mM NaCl. Washing buffer 2 contained 20 mM Tris-HCl, pH 7.4, 1.5 M NaCl, and 5% Triton X-100. Washing buffer 3 contained 20 mM Tris-HCl, pH 7.4, 500 mM NaCl, and 20 mM imidazole), and eluted by elution buffer (20 mM Tris-HCl, pH 7.4, 200 mM NaCl, and 250 mM imidazole). Then the proteins were dialyzed in dialysis buffer (20 mM Tris-HCl, pH 7.4, 50 mM NaCl, and 1 mM DTT). The purity of proteins was confirmed by SDS-PAGE and Coomassie blue staining, and concentrations were determined by a Coomassie Plus Protein Assay (Thermo Scientific).

### FRET measurement

We measured four sections of protein-protein interactions. First, recombinant CyPetRanGAP1c and YPetUbc9 proteins were incubated and mixed at room temperature in the Tris buffer (20 mM Tris-HCl, pH 7.5, 50 mM NaCl, DTT 1 mM) to a total volume of 60 μL. The final concentrations of CyPetRanGAP1c were fixed at 0.05, 0.1, 0.5, 1.0 μM and the final concentrations of YPetUbc9 were varied from 0 to 4 μM. Second, contaminating proteins which were extracted from pure BL21(DE3) E.coli cells without plasmid in it were added to CyPetRanGAP1c and YPetUbc9 proteins mixture. The concentrations of contaminating proteins were 1, 3, 10 μg in the Tris buffer (20 mM Tris-HCl, pH 7.5, 50 mM NaCl, DTT 1 mM) to a total volume of 60 μL. The final concentrations of CyPetRanGAP1c were fixed at 0.1, 0.5, 1 μM and the final concentrations of YPetUbc9 were varied from 0 to 4 μM. Third, pure BSA was added to CyPetRanGAP1c and YPetUbc9 proteins mixture. The concentration of BSA was 1 μg in the Tris buffer (20 mM Tris-HCl, pH 7.5, 50 mM NaCl, DTT 1 mM) to a total volume of 60 μL. The final concentrations of CyPetRanGAP1c were fixed at 0.05, 0.1, 0.5, 1.0 μM and the final concentrations of YPetUbc9 were varied from 0 to 4 μM. Forth, CyPetRanGAP1c and YPetUbc9 were extracted from BL21(DE3) E.coli cell without purification. We then measured the concentrations of CyPetRanGAP1c and YPetUbc9 by Fluorescence standard curve. The final concentrations of CyPetRanGAP1c were fixed at 0.05, 0.1, 0.5, 1.0 μM and the final concentrations of YPetUbc9 were varied from 0 to 4 μM. These sections of reaction were determined after mixing thoroughly, and the mixtures were examined with a fluorescence multi-well plate reader FlexstationII384 (Molecular Devices, Sunnyvale, CA). The fluorescence emission signals at 475 and 530 nm were collected under the excitation wavelength at 414 nm with a cutoff filter at 455 nm. Another fluorescence emission signals at 530 nm were collected at the excitation wavelength at 495 nm with a cutoff filter at 515 nm. The experiments were repeated three times, and the average values of fluorescence were recorded under each specific condition.

### Standard curve for CyPetRanGAP1c and YPetUbc9

The recombinant protein CyPetRanGAP1c and YPetUbc9 were incubated at 37°C in Tris buffer (20 mM Tris-HCl, pH 7.5, 50 mM NaCl, DTT 1 mM) to a total volume of 60 μL and added to each well of 384-well black/clear plate. The emission signals of CyPetRanGAP1c at 475 nm were collected after excitation at 414 nm, and the amount of protein was varied from 0 to 2 μM. The emission signals of YPetUbc9 at 530 nm were collected after excitation at 475 nm, and the amount of protein was ranging from 0 to 2 μM. When measuring the raw CyPetRanGAP1c and YPetUbc9 protein concentration, we used BL21 bacteria lysates as control.

### Fluorescence spectrum analysis of FRET

When the mixture was excited at 414 nm, emission peaks at 475 (FL_DD_) and at 530 nm (Em_total_) were obtained (see Fig. 2 B). When the mixture was excited at 495 nm, an emission peak at 530 nm (FL_AA_) was obtained. When a mixture of CyPetRanGAP1c and YPetUbc9 recombinant proteins was excited at 414 nm, nearly all of the emission intensity at 475 nm was the direct emission of CyPetRanGAP1c after energy transfer to YPetUbc9, and the emission intensity at 530 nm consisted of three components: the direct emission of CyPetRanGAP1c, the sensitized emission of YPetUbc9 (Em_FRET_) and the direct emission of YPetUbc9. Because Em_FRET_ is proportional to the amount of YPetUbc9 bound to CyPetRanGAP1c, we therefore can derive the relationship between Em_FRET_ and the bound concentration of YPetUbc9.

To determine the absolute Em_FRET_, two series of experiments were conducted to identify the ratio constants *α* and *β*. The first series of experiment was to determine the ratio constant of *α*. A series of CyPetRanGAP1c was prepared at concentrations of 0.05, 0.1, 0.5, 1.0 μM. The emissions of CyPetRanGAP1c at 475 and 530 nm were determined when excited at 414 nm. Dividing the fluorescence emission at 414 nm (FL_DD_) by its emission at 530 nm yielded the ratio constant, *α*. This is an estimate of unquenched CyPetRanGAP1c to the total emission at 530 nm when excited at 414 nm. In the second series of experiment, a series of YPetUbc9 was prepared at concentrations of 0.2, 0.5, 1 and 2, 3, and 4 μM. The emissions of YPetUbc9 at 530 nm were determined when excited at 414 or 495 nm. Dividing the fluorescence emission at 530 nm when excited at 414 nm by its emission at 530 nm when excited at 495 nm (FL_AA_) yielded the ratio constant, *β*.

### Data processing and *K_d_* determination

After the intensity of Em_FRET_ at each specific condition was calculated as described above, the dataset of Em_FRET_ and total concentration of YPetUbc9 ([YPetUbc9]_total_) was fitted by Prism 5 (GraphPad Software) to derive values for Em_FRETmax_ and *K_d_*. More specifically, the values of [YPetUbc9]_total_ were put into X-series, and the intensities of Em_FRET_ that were determined in triplicate at each [YPetUbc9]_total_ were put into Y-series. Nonlinear regression method was selected and a customized equation was created to fit the dataset:

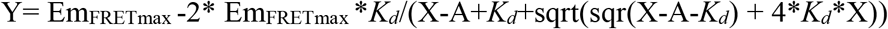

An equal to the concentration we added to the system, i.e., 0.05, 0.1, 0.5, or 1.0 μM. The initial values for the parameters Em_FRETmax_ and *K_d_* are set at 1.0, and one default constraint is that Em_FRETmax_ value must be greater than 0. The results were reported as mean ± SD.

### *K_d_* determination of the non-covalent RanGAP1c and Ubc9 interaction by SPR

His-tagged YPetUbc9 and CyPetRanGAP1c or His-tagged ubc9 and RanGAP1c dialysis stay overnight in running buffer (10 mM HEPES, 150 mM NaCl, 50 μM EDTA, 0.005% Tween20 pH7.4) to make sure all running conditions were the same. All analyses of interaction between CyPetRanGAP1c and YPetUbc9 or RanGAP1c and Ubc9 were performed on BIAcore X100 system equipped with NTA sensor chips (BIAcore AB, Uppsala, Sweden) at a flow rate of 30 μL/min. For immobilization of proteins, the chip was treated with 500 μM NiCl_2_ in running buffer for 1 min before 100 ng/mL purified YPetUbc9 or 200 ng/mL purified Ubc9 protein was injected for 120 s and stabilized for 120 s. Then 50~160 μg/mL thrombin-digested CyPetRanGAP1c or 10~40 μg/mL thrombin-digested RanGAP1c protein was injected for 120 s and disassociated for 10 min. To continuously monitor the nonspecific background binding of samples to the NTA surface, CyPetRanGAP1c and RanGAP1c proteins were injected into a control flow cell without treatment of NiCl_2_ and YPetUbc9/Ubc9 proteins. After determining one concentration of CyPetRanGAP1c and YPetRanGAP1c or RanGAP1c and RanGAP1c, the NTA sensor chip by regeneration buffer (10 mM HEPES, 150 mM NaCl, 350 mM EDTA, 0.005% Tween20 pH8.3) was regenerated, and then retreated by NiCl_2_, immobilized by YPetUbc9 or Ubc9. All measurements were performed at 25°C in running buffer. Data were analyzed with BIAcore X100 evaluation software ver.1.0 (BIAcore).

#### Statistical Analysis

All the Kd data under each combination of donor concentration and assay condition are reported as the mean ± standard error (SEM). The two-variable design offers the advantages of higher experimental efficiency and capability of statistically analyzing the interaction of the two factors. Two-way ANOVA test provides a statistical analysis to determine how the K_d_ values are affected by the two factors. Here, we implemented two-way ANOVA analysis using GraphPad Prism to compare the K_d_ values under different combination of donor concentration (row factor) and assay condition (column factor). This analysis divides the total variability among K_d_ values into four components: interactions between row and column, variability among columns, variability among rows, and variability among replicates. And it computes three P values that test three null hypotheses:

- Interaction P value: the null hypothesis is that there is no interaction between columns and rows.
- Column factor P value: the null hypothesis is that the mean of each column is the same.
- Row factor P value: the null hypothesis is that the mean of each row is the same

## RESULTS

### Theory of <u>H</u>igh-throughput determination of <u>p</u>rotein interaction <u>a</u>ffinity, *K_d_*, from <u>m</u>ixtures using quantitative <u>F</u>RET analysis (HPAMF)

To determine protein interaction dissociation constant, *K_d_*, we have developed a quantitative FRET analysis method to quantify FRET signal resulted from the protein interactive complex to elucidate the *K_d_* from either acceptor emission or donor quenching(Jiang *et al*, 2019; Song *et al*., 2011; Song *et al*, 2012). But we never explored this method to determine protein interaction affinity in the presence of other proteins or even from crude cell extracts (Fig.1A, left) using the FRET signal analysis method to differentiate fluorescence signals from each of free ligand, acceptor and ligand-acceptor complex (Fig.1A, middle), and Mass action principle (Fig.1A, right). This is not just a simple extension of the previous approach, but move this methodology into a totally new domain of protein interaction affinity determinations for difficult-to-be expressed or insoluble proteins. To achieve this, we first to determine the absolute FRET signals of interactive complex at each concentration and predict the maximum bound proteins from the titration experiment in the presence of other non-related proteins or cell extracts. We then could extend this methodology to determine FRET signals from two cell extracts containing each of fluorescence-labelled proteins as long as we can determine the two protein concentrations, which can be measured using external standard curves of two fluorescent protein signals.

**Figure 1.**
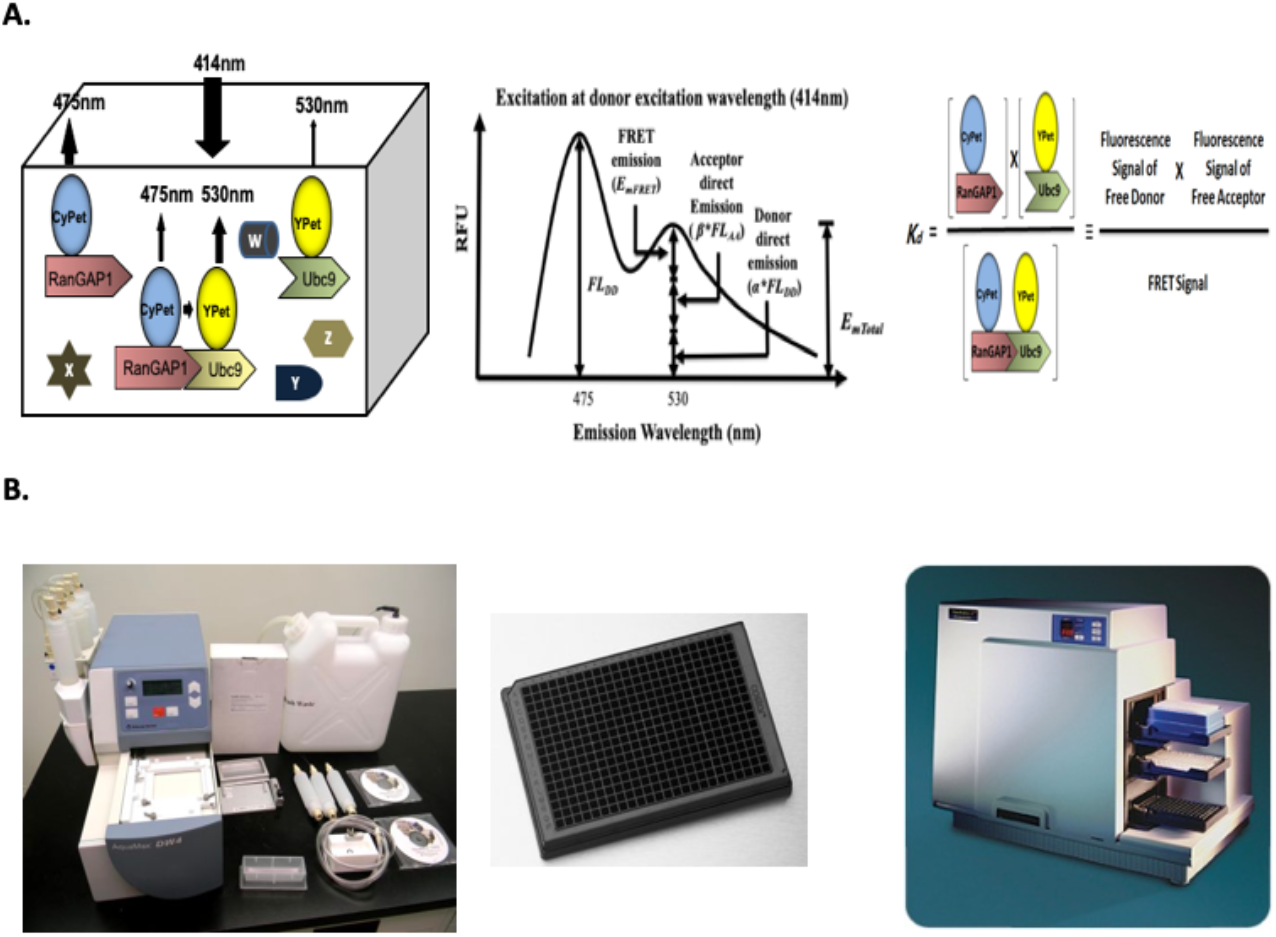
High-throughput FRET-based technology for protein interaction dissociation constant, *K_d_*, determination without purification. A. Schematic graph of fluorescence excitation and emission signals of interactive proteins, CyPet-RanGAPs and YPet-Ubc9, in the presence of other proteins(Left); quantitative FRET(qFRET) signal determination(Middle); and protein interaction dissociation constant, ***K_d_***, determination based on fluorescent signals of free donor and acceptor as well as interaction complex. B. High-throughput FRET assay for semi-automatic fluorescent protein distribution(Left); 384-well FRET assay platform(Middle); and Fluorescence plate reader, Flexstation^384^.

**Figure 2.**
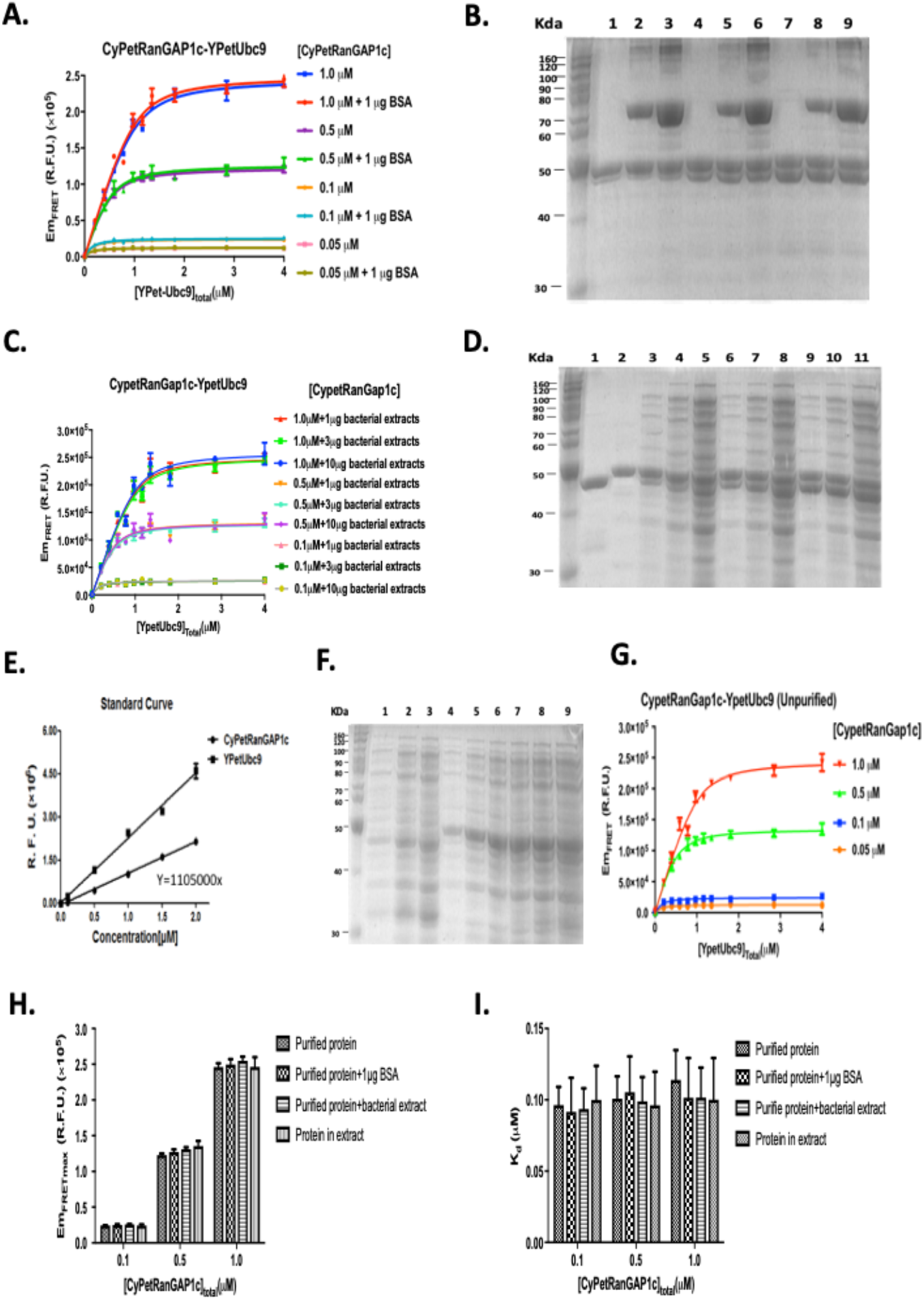
Determine CyPet-RanGAP1c and YPet-Ubc9 interaction affinity *K_d_* in the presence of BSA, or bacterial cell extract or from both extracts. **A.** Em_FRET_ determinations at concentrations of 0.05, 0.1, 0.5, and 1.0 μM of purified CyPetRanGAP1c, respectively, with increasing concentration of purified YPetUbc9 in the presence of 1mg BSA. **B.** The SDS-PAGE protein gel of CyPetRanGAP1c and YPetUbc9 without or with BSA stained with Coomassie. 0.1 μM (lane 1, 2, 3), 0.5 μM (lane 4, 5, 6) or 1.0 μM CyPetRanGAP1c (lane 7, 8, 9) + 1 μM YPetUbc9 (lane 1, 4, 7), with 1 μg BSA (lane 2, 5, 8), or with 3 μg BSA (lane 3, 6, 9). **C.** Em_FRET_ determinations at concentrations of 0.1, 0.5 and 1.0 μM of purified CyPetRanGAP1c, respectively, with increasing concentration of purified YpetUbc9 in the presence of E.coli lysates. **D.** The SDS-PAGE protein gel of CyPetRanGAP1c and YPetUbc9 without or with E.coli cell extract stained with Coomassie. CyPetRanGAP1c (lane 1); YPetUbc9 (Lane 2); 0.1 μM (lane 3,4 and 5), 0.5 μM (lane 6,7 and 8), 1.0 μM (lane 9,10 and 11) of CyPetRanGAP1c + 1 μM YPetUbc9 with 1 μg (lane 3,6 and 9), 3 μg (lane 4,7 and 10), 10 μg (lane 5,8 and 11) of E.coli cell extract. **E.** Standard curve of purified CyPet-RanGap1c or YPet-Ubc9 protein concentrations with its fluorescence emission signal. **F.** SDS-PAGE gel of E.coli cell crude extracts from cells expressing CyPert-RanGap1c (0.1mM, 0.5mM and 1mM in Lane 1-3, respectively), Ypet-Ubc9 (0.1mM, 0.5mM and 1mM in Lane 4-6, respectively), and mixture of CyPert-RanGap1c and Ypet-Ubc9 (0.1mM, 0.5mM and 1mM in Lane 7-9, respectively). **G.** Em_FRET_ determinations of CyPet-RanGAP1c at concentrations of 0.05, 0.1, 0.5 and 1.0 μM with increasing concentrations of YPet-Ubc9 from bacterial extracts. **H.** Maximum Em_FRET_ values of three concentrations of CyPetRanGAP1c with YPetUbc9 from purified proteins, or in the presence of BSA or bacterial extract, or both from bacterial cell extracts. **I.***K_d_* values from purified proteins, or in the presence of BSA or bacterial extract, or both from bacterial cell extracts.

To determine protein interaction *K_d_* from FRET signal, the first step is to obtain the absolute FRET signal (Em_FRET_) from the interactive protein complex in order to determine the maximum bound protein partner regardless for purified proteins in the presence of other proteins, or proteins in crude cell extracts. The direct emissions of the donor (CyPet) and acceptor (YPet) need to be determined and be excluded from the total emission at the FRET emission signal wavelength (530 nm in the case of the FRET pair, CyPet and YPet). However, the FRET emission signal wavelength at 530 nm consists of three components: the direct emission of CyPet, the direct emission of YPet, which do not come from the protein interaction, and the emission of the FRET signal (Em_FRET_), which comes from the protein interaction (Fig. 1A, middle). In order to obtain the Em_FRET_, the direct emissions of donor and acceptor need to be determined first. To measure the direct emissions in our FRET assay, we take a cross-wavelength coefficiency approach. First, the mixture of CyPetRanGAP1c and YPetUbc9 is excited at the donor excitation wavelength (414 nm), and two emission signals at 475 nm (FL_DD_) and 530 nm (FL_DA_) are determined, respectively. The fluorescence emissions at 475 nm when excited at 414 nm are from both the emission of CyPet (FL_DD_) and the direct emission of YPet at 475 nm, which is very small as compared with CyPet emission (<2.6% of CyPet emission) and can be neglected (unpublished data). We note that the direct emission of the donor CyPet at 530 nm is proportional to its emission at 475 nm when excited at 414 nm with a ratio factor of *α* (***α*\***FL_DD_), and the direct emission of YPet at 530 nm is proportional to its emission at 530 nm when excited at 495 nm with a ratio factor of *β* (*β*\*FL_AA_, FL_AA_ is the fluorescence emission of YPet at 530 nm when excited at acceptor excitation wavelength, 495 nm). After the emissions of FRET donor, CyPetRanGAP1c, at 475 nm when excited at 414 nm and FRET acceptor, YPetUbc9, at 530 nm when excited at 495 nm are determined, the FRET emission signal of the interactive complex, CyPetRanGAP1c/YPetUbc9, can be calculated by subtracting the above two signals with the ratios of *α* and *β*, respectively, from the total emission at 530 nm. In other words, the FRET emission signal (Em_FRET_) can be determined using the following formula:

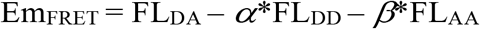

where the ratio constants *α* and *β* are first experimentally determined as 0.334±0.003 and 0.014±0.002, using free CyPetRanGAP1c and YPetUbc9 in this case, respectively.

Once we obtain Em_FRET_, following the general Mass action law,

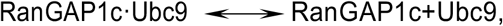

the dissociation constant, *K_d_*, can be defined as (Fig.1 B).

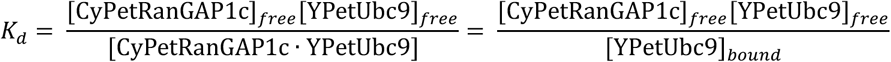

This can be rearranged to

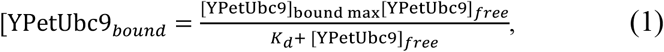

where [YPetUbc9]_boundmax_ is the theoretical maximal YPetUbc9 concentration that is bound to CyPetRanGAP1c concentration, and [YPetUbc9]_free_ is free YPetUbc9 concentration. [YPetUbc9]_bound_ is proportional to the FRET signal of bound proteins: Then Eq.(1) can be converted into Eq.(2) using the relationship,

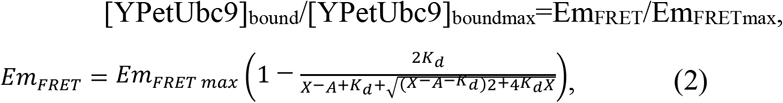

where Em_FRET_ is the absolute FRET signal and Em_FRET max_ is the absolute FRET signal when the maximum amount of YPetUbc9 is bound by CyPetRanGAP1c. A is the total concentration of CyPetRanGAP1c ([CyPetRanGAP1c]_total_), X as the concentrations of total YPetUbc9 ([YPetUbc9]_total_). Using this equation, the K_d_ and Em_FRET max_ can be determined.

The quantitative FRET assay has been carried out in the 384-well plate and fluorescence signals were determined using FlexStaion^384^ (Fig.1 B). Therefore, this approach can be implemented in a high-throughput technology platform that can be implemented for large-scale measurement.

### Determine interaction affinity *K_d_* of purified Ubc9 and RanGAP1c using HPAMF

We first determined the Em_FRET_ and *K_d_* of purified Ubc9 and RanGAP1c in order to establish the reference parameters of the interaction for later methodology development. In order to test the generality of the method, the CyPetRanGAP1c was set up in four concentrations, 0.05 μM, 0.1 μM, 0.5 μM and 1.0 μM accordingly, and the concentration of YPetUbc9 was gradually increased from 0 to 4 μM. The binding of YPetUbc9 to CyPetRanGAP1c resulted in an energy transfer from CyPet to YPet, and the emission at 530 nm was significantly increased, while the direct emission of CyPetRanGAP1c at 475 nm was decreased. We then determined the fluorescence emissions of each component in four different concentrations of CyPetRanGAP1c. The fluorescence emission spectra of all mixtures were then determined at the excitation wavelengths of 414 and 495 nm to exclude the direct emissions of CyPetRanGAP1c and YPetUbc9. After subtracting the direct emissions of CyPetRanGAP1c (***α*\***FL_DD_) and YPetUbc9 (***β*\***FL_AA_) at 530 nm from the total emission at 530 nm (FL_DA_), the absolute FRET emission at 530 nm when excited at 414 nm (Em_FRET_) increased steadily when more YPetUbc9 was added at each concentration of CyPetRanGAP1c. The estimated values for Em_FRETmax_, were (1.23±0.02)x10^4^, (2.43±0.04)x10^4^, (12.29±0.23)x10^4^, and (24.61±0.53)x10^4^, for 0.05, 0.1, 0.5, 1.0 μM of CyPetRanGAP1c, respectively(Supplement Fig.1 A) (Supplement Table 1). In this concentration range of the binding partner, the Em_FRETmax_ had a linear relationship with CyPetRanGAP1c from 3 to 60 pmole (Supplement Fig. 1B). This result suggests that our approach accurately and consistently predicted Em_FRETmax_ at various concentration ratios of CyPetRanGAP1c and YPetUbc9.

The disassociation constants of CyPetRanGAP1c and YPetUbc9 were then determined from the non-linear regression. By plotting Eq.(2) with Em_FRET_ vs. [YPetUbc9]_total_ in the Prism 5 program, we determined *K_d_*s from four concentrations of CyPetRanGAP1c (0.05, 0.1, 0.5, and 1.0 μM) as 0.098±0.014, 0.096±0.013, 0.101±0.016, and 0.114±0.021 μM, respectively, which were very close to the *K_d_* value determined using SPR,, 0.097μM, (Supplement Table 1) (Table 1). The very close values of *K_d_* generated from different concentrations of CyPet-RanGAP1c(from 0.05 μM to 1.0 μM of CyPetRanGAP1c) and various binding partner ratios of CyPetRanGAP1c to YPetUbc9 (from 0.67 to 40 folds) demonstrate that this method is very robust.

**Table 1.** Summary of *K_d_* values of CyPet-Gap1c and Ypet-Ubc9 from various conditions and as control, Surface Resonance Plasmon.

| | *K<sub>d</sub>*(μM) |
| --- | --- |
| **CyPetRanGAP1c-YPetUbc9** | **0.102±0.008** |
| **CyPetRanGAP1c-YPetUbc9+BSA** | **0.099±0.006** |
| **CyPetRanGAP1c-YPetUbc9+E.coli lysates** | **0.098±0.007** |
| **Crude CyPetRanGAP1c-YPetUbc9** | **0.100±0.003** |
| **Biacore™ SPR Systems** | **0.097** |

### Em_FRETmax_ and *K_d_* determinations in the presence of other protein(s) and molecules

To test our hypothesis that the **HPAMF** can be used for measuring *K_d_* in the presence of the other proteins and even total cell extract, we first determined the *K_d_* value in the presence of BSA and then bacterial cell extract. To show the robustness of the approach, we again conducted the assay in four concentrations of FRET donor CyPetRanGAP1c, 0.05 μM, 0.1 μM, 0.5 μM and 1.0 μM, and the concentration of YPetUbc9 was increased from 0 to 4 μM with an addition of 1ug of BSA, respectively. The Em_FRET_ were determined in the presence of BSA (Fig.2A). The CyPetRanGAP1c and YPetUbc9 were purified from bacterial cells using nickel agarose affinity chromatography. The purified proteins and added BSA were shown in SDS-PAGE gel (Figure 2B). From the gel, 0.1 μM(lane 1, 2, 3), 0.5 μM (lane 4, 5, 6) or 1.0 μM CyPetRanGAP1c (lane 7, 8, 9) + 1 μM YPetUbc9 (lane 1, 4, 7), with 1 μg BSA (lane 2, 5, 8), or with 3 μg BSA (lane 3, 6, 9). (Fig.2B).

The values for Em_FRETmax_, of the mixture in the presence of 1μg BSA were (1.26±0.04)x10^4^, (2.52±0.08)x10^4^, (12.71±0.38)x10^4^ and (24.97±0.76)x10^4^, for concentrations of 0.05, 0.1, 0.5, 1.0 μM of CyPetRanGAP1c, respectively(Supplement Table 1). By plotting Eq.(2) with Em_FRET_ and [YPetUbc9]_total_ in the Prism 5 program, we determined the *K_d_*s from four concentrations of CyPetRanGAP1c (0.05, 0.1, 0.5, 1.0) as 0.098±0.022, 0.092±0.024, 0.105±0.025, 0.102±0.028 μM, respectively(Supplement Table 1). The average *K_d_* is 0.099±0.006 μM. Comparing to the *K_d_* without BSA (0.102±0.008 μM), there is very close and no statistically significant difference (Table 1).

The disassociation constants of CyPetRanGAP1c and YPetUbc9 in the presence of bacterial extract were then determined in three concentrations of CyPetRanGAP1c (Fig.2C). From the gel in Fig. 2D, lane 1, 2 are pure CyPetRanGAP1c and YPetUbc9, 3~11 are pure CyPetRanGAP1c and YPetUbc9 mixture with 1 μg, 3 μg or 10 μg bacterial lysates proteins, respectively (Fig.2 D).

The values for Em_FRETmax_ of the mixture in the presence of E.coli lysates were 2.58±0.074, 13.26±0.43, and 25.21±0.90 when 1 μg E.coli lysates were added, respectively; 2.57±0.079, 13.02±0.39, and 25.19±0.99 when 3 μg E.coli lysates were added, respectively; 2.64±0.106, 13.07±0.58, and 26.06±1.14 when 10 μg E.coli lysates were added, respectively (Supplement Table 1).

When the concentration of CyPetRanGAP1c was fixed as 0.1 μM and 1, 3, 10 μg of bacterial extracts were added to the mixture, the values of *K_d_* were 0.092±0.022, 0.092±0.023, 0.096±0.031 μM, respectively (Supplement Table 1). When the concentration of CyPetRanGAP1c was fixed as 0.5 μM and 1, 3, 10 μg bacterial contaminated proteins were added to the mixture, the values of *K_d_* were 0.100±0.027, 0.108±0.025, 0.090±0.035 μM, respectively (Supplement Table 1). When the concentration of CyPetRanGAP1c was fixed as 1.0 μM and 1, 3, 10 μg bacterial contaminated proteins were added to the mixture, the *K_d_*s were 0.093±0.031, 0.109±0.037, 0.103±0.040 μM, respectively. Under these three conditions, the values of *K_d_* were very stable (Supplement Table 1). These results suggest that we can calculate *K_d_* by the quantitative FRET assay, and *K_d_*s determined in the absence or presence of BSA and bacterial extract using the quantitative FRET assay are very consistent.

So far, we have tried three different conditions, one with no contaminating protein, one with single contaminating protein, and another with multiple contaminating proteins. All these conditions gave us similar *K_d_* and Em_FRETmax_ values (Table 1). It indicates that our FRET-based method can be used to determine *K_d_* under complicated conditions. These values of *K_d_* determined by the quantitative FRET methods is close to the *K_d_* determined using the SPR described above(Table 1). This demonstrates that the **HPAMF** is a powerful approach for protein interactions and *K_d_* measurements.

### Determine interaction affinity *K_d_* of Ubc9 and RanGap1c directly from bacterial extracts

We then asked whether we can apply the HPAMF approach to determine the protein interaction Kd value directly from two cell extract without any purification. To achieve this, we first need to know the concentrations of the two FRET pair proteins in the cell extracts. The real concentrations of CyPetRanGAP1c and YPetUbc9 in the crude extracts were determined by monitoring the fluorescence signal at 475 nm and 530 nm from standard curve, respectively (Fig. 2E). The SDS-PAGE gel of the two bacterial cell extracts containing CyPetRanGAP1c and YPetUbc9 proteins, respectively, shown various proteins (Fig. 2F). YPetUbc9 was relatively easier to differentiate because its expression level is higher, while CyPetRanGAP1c is difficult to visualize the band in the gel. The concentrations of CyPetRanGAP1c was fixed at 0.05, 0.1, 0.5, 1.0 μM, respectively, and the concentrations of YPetUbc9 were increased from 0 to 4 μM. The Em_FRETmax_ was still linear as (1.308±0.041)x10^4^, (2.447±0.075)x10^4^, (13.57±0.39)x10^4^ and (24.63±0.79)x10^4^ (Supplement Table 1). Both Em_FRET max_ and *K_d_*s are very similar to the results from pure proteins interaction. The *K_d_*s were 0.102±0.024, 0.100±0.024, 0.096±0.023 and 0.100±0.029 μM, respectively. This result is surprisingly stable and the average of *K_d_* value, 0.100±0.003, agrees with the result from the pure proteins (Table1).

Further analysis shows no statistically significant difference among these Em_FRETmax_ and *K_d_* values in all the four conditions, demonstrating the reliability and robustness of the FRET-based *K_d_* determination approach using un-purified proteins (Fig. 2H and I). This demonstrate the HPAMF method is very reliable and robustness.

### HPAMF shows high affinities of SUMOylation E3 ligase with the E2 conjugating enzyme

To test its ability to determine protein interaction affinity that is generally regarded as very challenging to determine, we applied the HPAMF to determine the affinity of SUMOylation E3 ligase, PIAS1, and E2 conjugating enzyme, Ubc9. SUMOs (SUMO1-4) are ubiquitin-like polypeptides that are covalently conjugated to target proteins as one type of major protein post-translational modifications [2-4]. SUMO conjugation occurs through an enzymatic cascade, involving E1-activating enzyme, E2-conjugating enzyme and E3 protein ligases[5]. *In vitro*, with the help of E1 enzyme and E2 enzyme, Ubc9 can recognize and transfer SUMO peptide to most substrates directly, whereas the E3 ligases, such as PIAS1, is speculated to facilitate most SUMOylation in cells by recruiting E2~SUMO and substrate into a complex to promote SUMO conjugation *in vivo*. and stimulating SUMO conjugate to lysine residues of substrates under physiological conditions[6-10]. Although genetics and non-quantitative biochemical experiments have shown that PIAS1 is E3 ligase for SUMOylation, and the PIAS1, as SUMO E3, has never been shown in kinetics to interact with Ubc9 at high affinity so far, partially due to its difficulty to be expressed in many expression systems.

We therefore applied the HPAMF to determine the affinity of PIAS1 and Ubc9 interaction. We cloned the PIAS1 cDNA after codon optimization, and Ubiquitin E2 enzyme the Ubc12 cDNA as control, as fusion proteins with CyPet and YPet, respectively, into pET28(b) vector. We optimized the codons of PIAS1 because almost no PIAS1 protein could be expressed and purified with its original codons. The plasmids were transfected into E.coli Bl21(DE3) cells and induced overnight with IPTG. The protein expressions were examined with SDS-PAGE gel for samples of uninduced and induced cells, and supernants after centrifugation of sonicated cells (Fig, 3A). The CyPet-Ubc9 and CyPet-Ubc12 were well induced (Figure 4A, lane1-3 and 7-9), while the YPet-PIAS1 protein was hardly seen (Figure 4, lane 4-6). We then calculated the CyPet-Ubc9 and CyPet-Ubc12, and YPet-PIAS1 amounts using external CyPet and YPet standard curves, respectively. We then performed the FRET titration experiments. In each titration experiment, the concentration of CyPet-Ubc9 or Cypet-Ubc12 was fixed and increasing concentrations of YPet-PIAS1 were added. The values of Em_FRERT_ in each point were calculated according to Eq.(2). The titration curves of Em_FRERT_ are shown (Fig. 3B). The titration curves of CyPet-Ubc9 and YPet-PIAS1 shown good dose-dependent increase with the concentrations of CyPet-Ubc9, while the titration curves of CyPet-Ubc12 with YPet-PIAS1 did not show any Em_FRERT_ signal even at the highest concentration of YPet-PIAS1, suggesting a strong specific interaction between Ubc9 and PIAS1. We then determined the interaction affinities of CyPet-Ubc9 with YPet-PIAS1 at each concentration of CyPet-Ubc9 (Figure 4C). The *K_d_* values were very consistent, ranging from 0.26 ± 0.04 to 0.30 ± 0.04 from the four testing conditions. This result suggests that PIAS1 interacts with Ubc9 at very high affinity, and therefore, set up the basis of specificity. Further analysis shows no statistically significant difference in these *K_d_* values, again demonstrating the reliability of the HPAMF.

**Figure 3.**
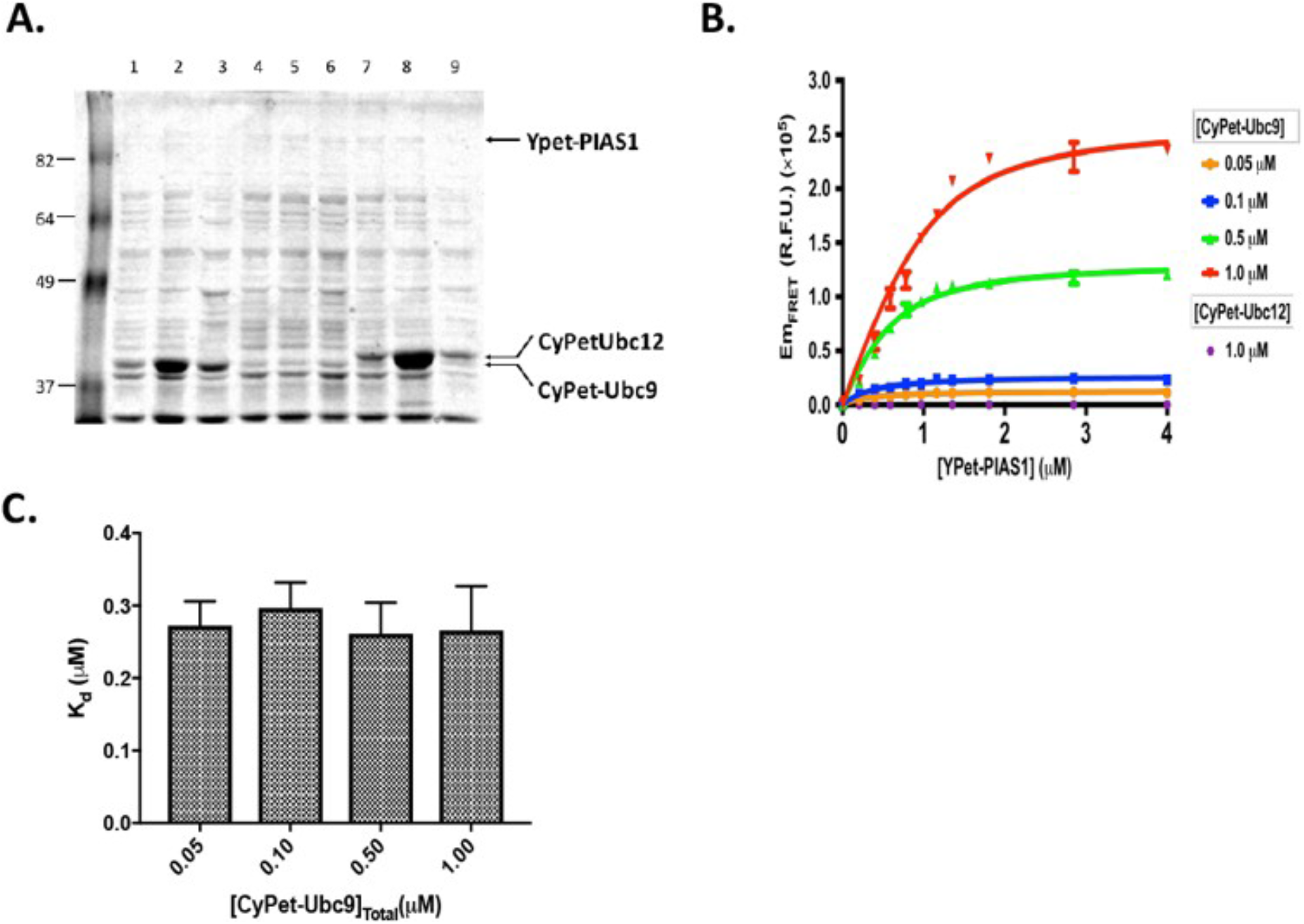
Determine specificity of SUMO E3 ligase, PIAS1, to its E2 conjugating enzyme, Ubc9, from interaction affinity. **A.** SDS-PAGE gel of E.coli cells of un-induced (lane 1, 4,and 7), induced (lane 2,5,and 8) or supernant (lane 3,6, and 9) of CyPet-Ubc9, YPet-PIAS1, CyPet-Ubc12, respectively. **B.** Em_FRET_ determinations of various concentrations of CyPet-Ubc9 or CyPet-Ubc12 with increasing concentrations of YPet-PIAS1. **C.** *K_d_* determinations of PIAS1 with Ubc9 from various concentrations of CyPet-Ubc9.

**Figure 4.**
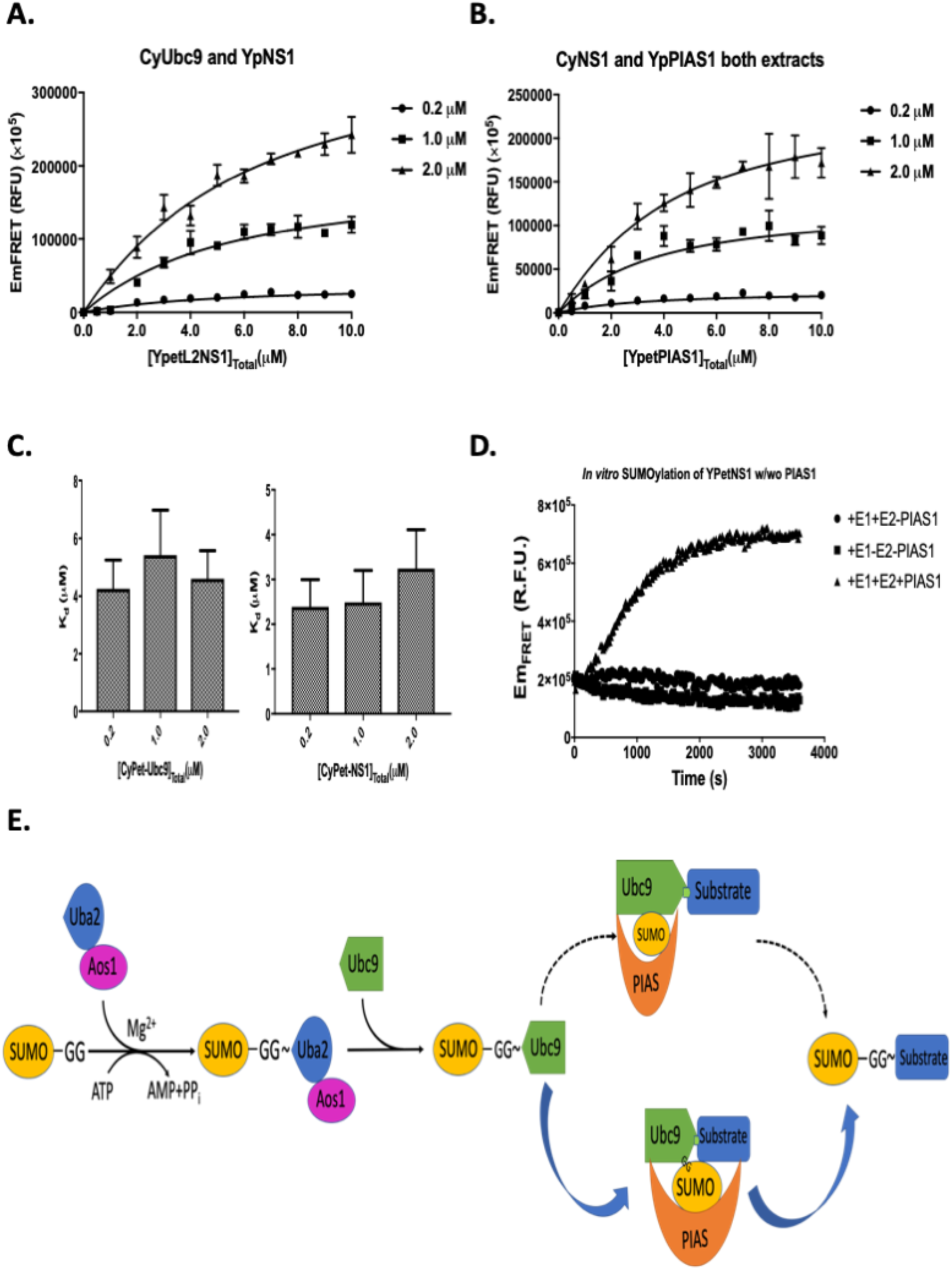
Both SUMO E2 conjugating enzyme E2 and E3 ligase contribute to substrate recognition. **A.** E2 Ubc9 recognizes influenza virus protein NS1 at high affinity. **B.** E3 PIAS1 recognizes influenza virus protein NS1 at high affinity. **C.** The bar graph of *K_d_* values of NS1 with Ubc9 and PIAS1, respectively. **D.** PIAS1 facilitates the SUMOylation of NS1. E. The model of substrate recognition in the SUMOylation cascade.

### SUMO E3 ligase recognize influenza virus protein NS1 directly with high affinity

It has been suggested that there may be two possible models on how SUMOylation cascade can transfer the SUMO peptide to substrates: the first one was that both E2 and E3 would interact with substrates simultaneously; the second was that only E2 would interact with substrates, while E3 only brings E2 and SUMO peptide together to the substrates, and E3 does not recognize substrates. To distinguish these two models from the kinetics perspective, we used the HPAMF to determine the interaction affinities of Ubc9 and PIAS1 to SUMOylation substrate, influenza virus protein NS1, respectively. The CyPet-Ubc9 and YPet-NS1 were expressed and purified from bacterial Bl21(DE3) cells. The Em_FRET_ of CyPet-Ubc9 and YPet-NS1 was determined in three concentrations of CyPet-Ubc9 (Fig. 4A). Then we derived the interaction affinity of PIAS1 with NS1. The YPet-PIAS1 and CyPet-NS1 were expressed in bacterial cells and supernant mixtures were prepared after sonication. According to the CyPet and YPet standard curves, the concentrations of YPet-PIAS1 and CyPet-NS1 were determined. Then we determined the Em_FRET_ in three concentrations of CyPet-NS1, 0.2μM, 1.0μM and 2.0μM, respectively (Fig. 4B). The average *K_d_* value was 2.96 ± 0.47 uM, slightly higher than that of UBC9 to NS1, which average was 4.84 ± 0.64 μM (Figure 5C). This result suggests that, although both Ubc9 and PIAS1 interact with substrate at modest affinities, they work together to contribute to the substrate recognition, and therefore, conferring higher affinity.

To determine the contribution of SUMO E3 ligase PIAS1 during the SUMOylation process, we then carried out a full SUMOylation assay with or without PIAS1 using FRET assay in a different mode. In this setting, if the SUMOylation happened, the conjugation of CyPet-SUMO1 to YPet-NS1 from the SUMOylation cascade would lead to FRET signal generation. The SUMOylation assay was carried out with CyPet-SUMO1, Aos1/Uba2, Ubc9 and YPet-NS1 in the presence or absence of PIAS1. In this assay with low concentration of Ubc9(1uM), the SUMO conjugation only happened in the presence of PIAS1, and as a control, no FRET signal was observed without ATP (Fig. 4D). This result indicates that PIAS1 can help the SUMO conjugation and may reflect physiological conditions, in which PIAS1 has been discovered to be requited for some substrate SUMOylation *in vivo(Liu et al, 2004)*.

From these data, we propose the following mechanistic model of SUMOylation process (Fig.4E). After the activation of SUMO peptides by E1 and E2 enzymes, the SUMO peptides are conjugated to substrates with the substrate recognitions by both E2 and E3 enzymes, therefore, conferring high affinity and specificity. This model supported by kinetics parameters may explain the long-term mystery of SUMOylation specificity *in vivo*.

## DISCUSSION

In current study, we present a novel approach (HPAMF) using quantitative FRET analysis for protein interaction affinity determinations directly from cell extracts without any purification, and provide a kinetics basis for the mechanistic understanding of SUMOylation E2/E3 co-recognition of substrates *in vivo*. This provides a clear explanation of SUMOylation susbrtate specificity from a new angel of interaction affinity. Applications of our approach to various protein-protein interactions should provide high-quality protein interaction affinities of systems, network, proteomes, and provide comprehensive quantitative biological and biomedical maps without a need of laborious protein purification, especially for those difficult-to-be expressed proteins. Our work represents a large step forward toward enabling quantitative protein interactive network within an organism.

In this research, we also verified our results by the SPR method, which demonstrates that the novel quantitative FRET assay produces consistent results that agree well with those determined using classical *K_d_* measurement technologies. Compared with SPR, the FRET-based method is better to determine protein-protein interaction, as the HPAMF does not require protein purifications. The SPR may have disadvantages in some cases, such as proteins immobilized on sensor surface may not be their native conformation and heterogenicity of binding due to different orientation of ligand immobilized on the surface may generate decreased affinity(Helmerhorst *et al*, 2012; Schuck & Zhao, 2010). Also due to surface immobilization of ligand, the local concentration is higher than the solution, the binding kinetic is different from ideal pseudo-first-order binding due to mass transfer effect(Schuck & Minton, 1996; Schuck & Zhao, 2010). In addition, rebinding effect and nonspecific binding to the sensor chip may occur and thus, more careful mathematic algorithms are needed to obtain meaningful parameters(Nieba *et al*, 1996). This SPR method for *K_d_* determination might not be valid when a simple Langmuir-type binding model does not apply(O’Shannessy & Winzor, 1996; Schuck & Minton, 1996). In addition, one concentration of protein is used in the SPR assay in each time and it has to wait for 40 min to get the final result. So it won’t feasible for large sample testings. If some enzymes are involved in the final product formation, leading to the interaction, it may inconsistent results as enzymes may lost activities during this period of time. In contrast, six concentrations and three time repeats of interaction only took less than 1 hour in the HPAMF method. Also our HPAMF method has already taken into account the potential orientation problem as it determine interactions in solution. In addition to SPR, many different methods, which include Isothermal Titration Calorimetry (ITC), radioactive labeling and ultracentrifugation, have also been used for *K_d_* determination(Syafrizayanti *et al*., 2014; Zhou *et al*., 2016). These methods offer experimental convenience, but also have some disadvantages. They often require the environmentally unfriendly labeling or expensive instrumentation, and especially can’t determine impure proteins interactions. Isothermal titration calorimetry (ITC) requires relatively large amounts (i.e., micromolar range) of samples and systematics errors in cell volume, the heart calibration, and other issues, such as baseline errors and gas bubbles can lead to inaccuracies(Tellinghuisen, 2004, 2007; Tellinghuisen & Chodera, 2011). It also requires a relatively expensive dedicated equipment. The elongated centrifugation can perturb the equilibrium between bound and free proteins, especially if the dissociation rates are fast, and thus *K_d_* values determined may not represent true equilibrium constants. In ultracentrifugation assay, peripheral proteins can be nonspecifically adsorbed on the test tube walls during high-speed centrifugation. In principle, the fluorescent polarization (FP) approach can be used potentially to address these issues. It has been successfully used to determine protein interaction dissociation constant and has been developed into HTS platform for small molecule drug screening. However, the FP approach also needs purified interacting partners and is less sensitive as it only measures the polarized fluorescent signal(Park & Raines, 2004). It also typically requires very sensitive instruments for good quantification, such as fluorescent microscopy, preventing its high-throughput application.

Among the methods evaluated, our HPAMF approach is the only one capable of determining protein interaction affinity without purification and in high-throughput mode. The HPAMF also provides kinetics parameters of multi-protein reactions in protein cascade reactions. The additional ability to link protein interaction affinity with multi-protein reaction kinetics provides an opportunity to investigate biochemical reactions in a complex environment. Extension of qFRET approaches to biochemical reactions will probably allow generations of far more comprehensive quantitative map of biochemical reactions in life. Further applications of our HPAMF to diverse protein-protein interactions, especially those in which expression and purification have been limited, should help understand vast protein interactions that are current unknown and provide a universal measurement for quantitative systems biology.

The consistent affinity results of the *K_d_* determination at nanomolar range show that our method is not only accurate and reliable at various concentrations of the interactive partners but also sensitive at high affinity nanomolar level. In contrast, traditional radio-labeled protein binding assays to determine *K_d_* requires a range of at least 100-fold of labeled ligand to predict maximum binding. Our FRET-based *K_d_* determination approach accurately determines *K_d_* at ratios 0.67–40 fold of the binding partners of the RanGAP1c-Ubc9. Other approaches for the *K_d_* determination, such as SPR or isothermal titration calorimetry, require multiple steps and special instruments and often give large variations. While the FRET assay has become more popular in biochemical and cell biology studies, our quantitative FRET method would advance FRET technology to another quantitative level, and information on RanGAP1c-Ubc9 interaction affinity will provide valuable insights into the complex UBL conjugation cascade from systems biology perspective.

## ACKNOWLEDGEMENTS

We are very grateful to Dr. Michael Pirrung in the Department of Chemistry, University of California at Riverside for allowing us to access his molecular biology lab. We thank all the members in Liao’s group for their very close collaborative work and assistance with this study.

## FUNDING

The work was partially supported by the UCR Academic Senate Grant and Attaisina Gift Grant. Conflict of interest statement. None declared.

---

**Supplement Table 1.** Summary of the maximal FRET emission Em_FRETmax_ and *K_d_* values in different conditions.

| CyPetRanGAP1c(μM) | $K_d$(μM) | $Em_{FRETmax}$ (R.F.U.)($\times 10^4$) |
| --- | --- | --- |
| 0.05 | 0.098±0.014 | 1.227±0.022 |
| 0.1 | 0.096±0.013 | 2.433±0.041 |
| 0.5 | 0.101±0.016 | 12.29±0.23 |
| 1 | 0.114±0.021 | 24.61±0.53 |
| 0.05+1 μgBSA | 0.098±0.022 | 1.260±0.036 |
| 0.1+1μg BSA | 0.092±0.024 | 2.523±0.080 |
| 0.5+1 μg BSA | 0.105±0.025 | 12.71±0.38 |
| 1.0+1 μg BSA | 0.102±0.028 | 24.97±0.76 |
| 0.1+1μg bacterial extract | 0.092±0.022 | 2.583±0.074 |
| 0.1+3μg bacterial extract | 0.092±0.023 | 2.572±0.079 |
| 0.1+10μg bacterial extract | 0.096±0.031 | 2.636±0.106 |
| 0.5+1μg bacterial extract | 0.100±0.027 | 13.26±0.43 |
| 0.5+3μg bacterial extract | 0.108±0.025 | 13.02±0.39 |
| 0.5+10μg bacterial extract | 0.090±0.035 | 13.07±0.58 |
| 1.0+1μg bacterial extract | 0.093±0.031 | 25.21±0.90 |
| 1.0+3μg bacterial extract | 0.109±0.037 | 25.19±0.99 |
| 1.0+10μg bacterial extract | 0.103±0.040 | 26.06±1.14 |
| 0.05 (from crude extract) | 0.102±0.024 | 1.308±0.041 |
| 0.1 (from crude extract) | 0.100±0.024 | 2.447±0.075 |
| 0.5 (from crude extract) | 0.096±0.023 | 13.57±0.39 |
| 1.0 (from crude extract) | 0.100±0.029 | 24.63±0.79 |
| SPR Method *by Ling* | 0.097 |  |

**Supplement Figure 1.**
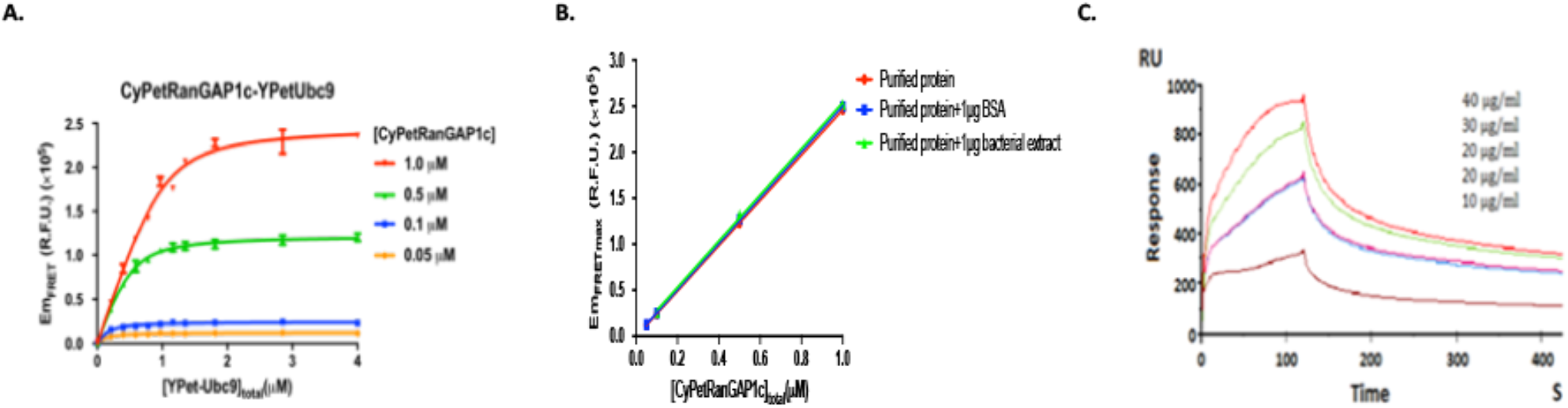
Interaction affinity determination of CyPet-Ubc9 and Ypet-PIAS1. **A.** Em_FRET_ determinations of purified CyPet-Ubc9 at concentrations of 0.05mM, 0.1mM, 0.5mM and 1.0mM with increasing concentrations of purified Ypet-PIAS1. B. Determinations of Em_FRETmax_ at different concentrations of the donor CyPet-RanGAP1C with YPet-Ubc9 in the absence and presence of other proteins. The maximal FRET emission is proportional to the amount of CyPet-RanGAP1 in the assay with or without other proteins. C. Dendrogram CyPet–RanGAP1c and YPet–Ubc9 interaction by the surface plasma resonance (SPR).

